# Meta-analysis of the melanoma miRNome identifies consensus signatures of progression and prognosis regulating metabolic plasticity and stress resistance

**DOI:** 10.1101/2023.10.20.563284

**Authors:** Borja Gómez-Cabañes, Helena Gómez-Martínez, Cristina Galiana-Roselló, Zoraida Andreu, Adolfo López-Cerdán, Nicole M. Eskow, José A López-Guerrero, Beatriz Dolader-Rabinad, Ricardo A Hernández-Ramírez, Irene Soler-Sáez, Marta R. Hidalgo, Eva Hernando, Francisco García-García

**Affiliations:** Computational Biomedicine Laboratory, Principe Felipe Research Center (CIPF). Eduardo Primo Yúfera 3, Valencia, 46012, Spain; Programa de Doctorado en Biotecnología, Universitat Politècnica de València. Camí de Vera s/n, 46022 València, Spain; Laboratory of Molecular Biology, Foundation of the Valencian Institute of Oncology, Valencia, 46009, Spain; IVO-CIPF Joint Cancer Research Unit, Príncipe Felipe Research Center (CIPF). Eduardo Primo Yúfera 3, Valencia, 46012, Spain; Joint unit in Artificial Intelligence and Biomedical Imaging FISABIO-CIPF, Foundation for the Promotion of Health and Biomedical Research of Valencia Region, Valencia, Spain; Department of Pathology, New York University Grossman School of Medicine, New York, US; Department of Pathology, Catholic University of Valencia, Valencia, Spain; Area of Applied Mathematics, Department of Applied Mathematics, Universidad de Valencia;Valencia, 46010, Spain

**Keywords:** Systematic review, meta-analysis, melanoma, diagnosis, microRNA

## Abstract

Cutaneous melanoma is the deadliest form of skin cancer, and accurate risk stratification remains challenging because of the molecular heterogeneity underlying disease progression. The transition from benign nevi to malignant melanoma and the development of the high-risk ulcerated phenotype require novel biomarkers to improve diagnosis and prognosis. We aimed to integrate miRNA-level evidence to generate consensus signatures associated with these processes. We performed a robust meta-analysis of six independent transcriptomic studies, overcoming inter-study heterogeneity, combined with extensive *in silico* characterization and validated miRNA–mRNA interactome construction. In the diagnostic setting, we identified a consensus signature of 24 miRNAs. Network topology analysis highlighted hsa-miR-142-5p as a candidate regulatory hub targeting CDK6, SIRT1, and TGFBR2, potentially contributing to the disruption of Oncogene-Induced Senescence barriers. In the prognostic setting, we identified a stress-adaptive signature of 23 miRNAs, with hsa-miR-223-3p emerging as a candidate driver of invasiveness through predicted RHOB suppression, while loss of hsa-miR-200c may facilitate epithelial–mesenchymal transition and EGFR-mediated survival pathways. Overall, our findings support a sequential model in which distinct miRNA alterations first promote senescence escape and later favor adaptation to the ulcerated microenvironment. These candidate regulatory axes warrant experimental validation as potential biomarkers and therapeutic targets.

**Key points:** *● Why was the study undertaken?:* The study was undertaken to address persistent challenges in cutaneous melanoma risk stratification, which remain difficult due to the molecular heterogeneity underlying disease progression. Specifically, there is an urgent clinical need to identify novel miRNA biomarkers to resolve critical barriers in disease management, particularly during the transition from benign nevi to malignant melanoma and the progression toward the high-risk ulcerated phenotype, both of which currently lack precise molecular determinants.

***●** What does this study add?:* By integrating dispersed transcriptomic data, we deliver a robust meta-analysis that identifies two consensus miRNA signatures. We define a diagnostic hub, hsa-miR-142-5p, linked to senescence escape, and a ’stress-adaptive’ prognostic signature for high-risk ulcerated phenotype. Our study adds a novel mechanistic layer to disease progression, specifically identifying the hsa-miR-223-3p/RHOB axis as a driver of invasiveness, supported by validated miRNA-mRNA interactomes that map the regulatory architecture of early melanomagenesis and later stress adaptation.

***●** What are the implications of this study for disease understanding and/or clinical care?:* Our findings propose a sequential model of progression: miRNA alterations initially facilitate senescence escape, followed by active stress adaptation in the ulcerated niche. This provides a multi-layered view of tumor plasticity, identifying candidate therapeutic targets such as CDK4/6 and EGFR signaling pathways. Clinically, these data generate testable experimental hypotheses, suggesting that targeting specific miRNA-driven regulatory axes, particularly those regulating metabolic plasticity and anoikis resistance, could overcome therapeutic resistance and improve risk stratification in aggressive melanoma.

## Introduction

Cutaneous melanoma represents the most lethal form of skin cancer, accounting for 80% of related mortalities despite comprising only 5% of cases (1,2). While the 5-year survival rate exceeds 98% for localized epidermal lesions (3), prognosis deteriorates precipitously once the tumor invades the dermis during the vertical growth phase (4). Crucially, histopathological ulceration serves as an independent T-stage modifier in the AJCC 8th edition staging system and a key predictor of poor outcome (5). However, the tumor’s intrinsic biological heterogeneity often hinders early detection (6), and the specific miRNA-level molecular determinants driving these aggressive transitions remain incompletely characterized, challenging precise risk stratification.

MicroRNAs (miRNAs) are short non-coding RNAs that regulate the expression of genes associated with numerous biological processes. The dysregulated expression profiles of miRNAs in tumor tissue make them promising biomarkers for cancer diagnosis and prognosis (7–12). Although individual studies have linked specific miRNAs to mutational status, proliferation, and survival outcomes, including in melanoma, (13–17), significant research gaps persist regarding the reproducibility of these findings across independent cohorts. This lack of consistency demands systematic cross-validation approaches to robustly connect miRNA alterations with established clinical outcome predictors.

To address these limitations, we conducted a systematic review and meta-analysis adhering to PRISMA guidelines (18), integrating datasets to present a consensus miRNA signature across both diagnostic and prognostic clinical scenarios. In the diagnostic context, we identified a core signature of 24 differentially expressed miRNAs driven by hsa-miR-1973 and hsa-miR-142-5p, exposing early molecular mechanisms of senescence escape. Concurrently, in the prognostic scenario, we established a 23-miRNA signature where the hsa-miR-223-3p/hsa-miR-200c axis emerges as a key indicator of the high-risk ulcerated phenotype, potentially orchestrating metabolic adaptation and anoikis resistance. This comprehensive *in silico* framework provides critical insights into the biological roles and clinical relevance of these miRNA markers.

## Results

### Systematic review and meta-analysis of miRNA studies

From 54 screened entries, six independent transcriptomic studies published between 2010 and 2023 (N = 303 skin tissue specimens) met all inclusion criteria (**Figure 1**). These datasets were stratified into a diagnostic scenario, which involved a comparison between primary cutaneous melanoma and benign nevi (PM vs. BN); and a prognostic scenario, which involved a comparison between ulcerated and non-ulcerated primary cutaneous melanoma (UM vs. NUM). The specific scenarios subjected to analysis, along with the corresponding number of differentially expressed miRNAs identified in each study, are summarized in **Table S1** and **Table S2**, respectively.

**Figure 1.**
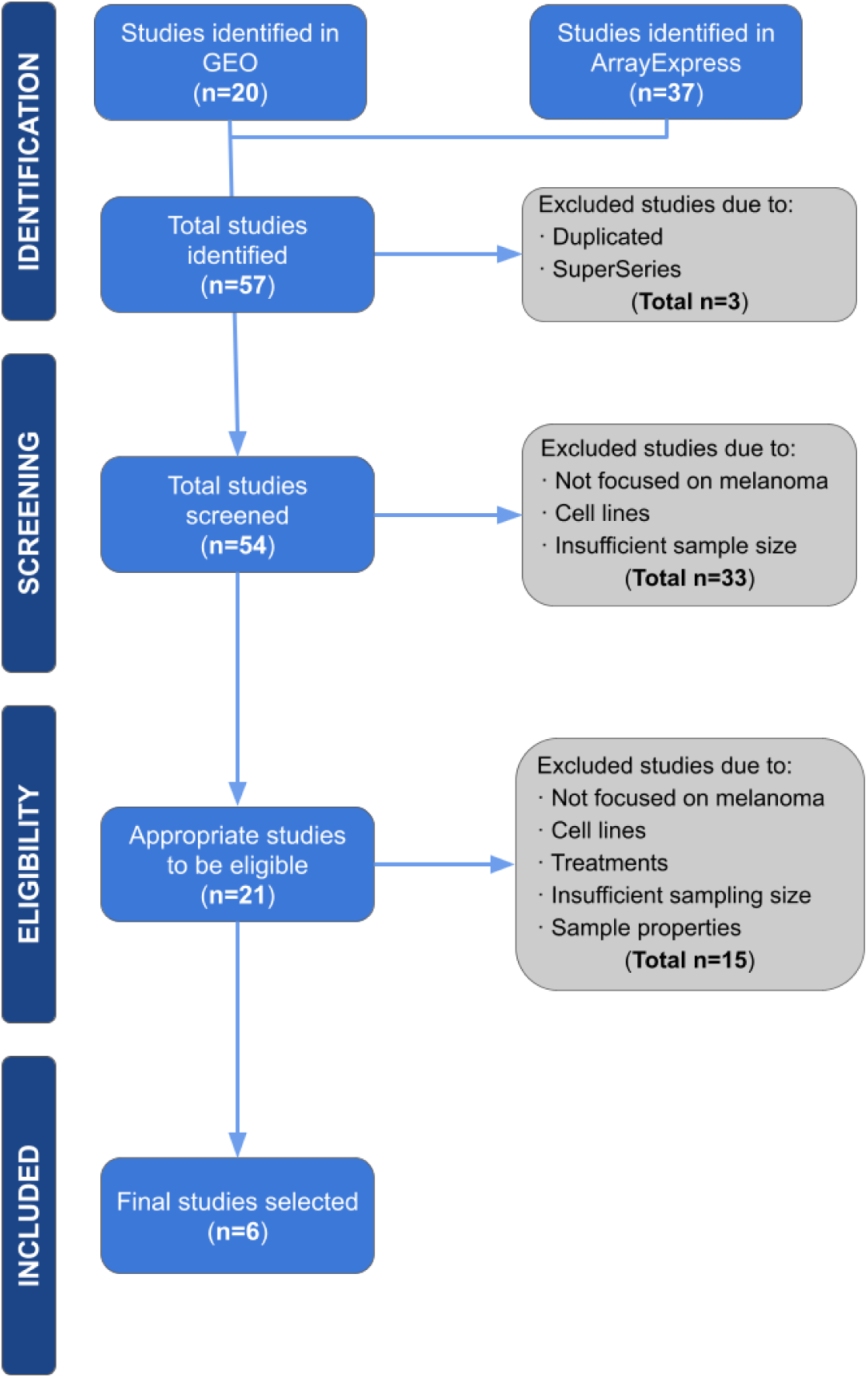
PRISMA diagram of the systematic review.

Random-effects meta-analysis revealed a consensus diagnostic signature of 24 differentially expressed miRNAs (11 upregulated and 13 downregulated in PM) and a prognostic signature of 23 miRNAs (7 upregulated and 16 downregulated in UM). Three miRNA markers overlapped across both clinical settings with identical directional patterns: hsa-miR-21 was consistently upregulated, while hsa-miR-203 and hsa-miR-205 were consistently downregulated. The complete results for significant miRNAs are presented in **Table 1** and **Table S3**.

**Table 1.** Significant differentially expressed miRNAs identified by random-effects meta-analysis in diagnostic and prognostic scenarios. Data represent the consensus signatures for the Primary Melanoma vs. Benign Nevi (Diagnostic) and Ulcerated vs. Non-Ulcerated Melanoma (prognostic) comparisons. For each miRNA, the combined log2 fold-change (logFC) and the adjusted P-value are reported. Selection criteria for significance were defined as an adjusted P-value < 0.05 and an absolute logFC > 0.5.

| Primary melanoma vs. Benign nevus |  |  | Ulcerated vs. Non-ulcerated |  |  |
| --- | --- | --- | --- | --- | --- |
| miRNA | logFC | adjusted P-value | miRNA | logFC | adjusted P-value |
| hsa-miR-1973 | 3.556 | 0.0002 | hsa-miR-223-3p | 1.494 | 0.0070 |
| hsa-miR-3648 | 3.340 | 0.0002 | hsa-miR-196b-5p | 1.351 | 0.0370 |
| hsa-miR-3156-5p | 3.072 | 0.0120 | hsa-miR-135b-5p | 1.262 | 0.0240 |
| hsa-miR-3911 | 1.926 | 0.0040 | hsa-miR-363-3p | 1.195 | 0.0210 |
| hsa-miR-21 | 1.776 | 0.0002 | hsa-miR-451 | 1.049 | 0.0001 |
| hsa-miR-3610 | 1.275 | 0.0380 | hsa-miR-21 | 0.788 | 0.0010 |
| hsa-miR-3138 | 1.274 | 0.0380 | hsa-miR-29a | 0.526 | 0.0390 |
| hsa-miR-3195 | 1.156 | 0.0150 | hsa-miR-200c | -0.501 | 0.0020 |
| hsa-miR-3132 | 0.999 | 0.0380 | hsa-miR-5010-5p | -0.511 | 0.0480 |
| hsa-miR-142-5p | 0.814 | 0.0020 | hsa-miR-301a-5p | -0.515 | 0.0240 |
| hsa-miR-22 | 0.754 | 0.0380 | hsa-miR-450a-2-3p | -0.539 | 0.0390 |
| hsa-miR-23a | -0.561 | 0.0380 | hsa-miR-580-3p | -0.542 | 0.0480 |
| hsa-miR-30c | -0.579 | 0.0190 | hsa-miR-381-5p | -0.545 | 0.0400 |
| hsa-let-7d | -0.632 | 0.0020 | hsa-miR-203 | -0.561 | 0.0190 |
| hsa-let-7c | -0.646 | 0.0380 | hsa-miR-487b-5p | -0.590 | 0.0480 |
| hsa-miR-193b | -0.747 | 0.0002 | hsa-miR-205 | -0.593 | 0.0190 |
| hsa-miR-508-3p | -1.040 | 0.0002 | hsa-miR-552-3p | -0.597 | 0.0390 |
| hsa-miR-509-3p | -1.305 | 0.0002 | hsa-miR-499a-5p | -0.633 | 0.0030 |
| hsa-miR-23b | -1.323 | 0.0002 | hsa-miR-518e-3p | -0.649 | 0.0480 |
| hsa-miR-203 | -1.368 | 0.0160 | hsa-miR-489-3p | -0.659 | 0.0480 |
| hsa-miR-205 | -1.571 | 0.0020 | hsa-miR-133a-5p | -0.667 | 0.0190 |
| hsa-miR-100 | -1.700 | 0.0002 | hsa-miR-624-3p | -0.669 | 0.0480 |
| hsa-miR-101 | -1.785 | 0.0002 | hsa-miR-376c-5p | -0.749 | 0.0260 |
| hsa-miR-211 | -1.956 | 0.0080 |  |  |  |

To validate the prognostic signature, an orthogonal cross-validation was conducted against an independent Stage II/III melanoma cohort from the InterMEL consortium profiled via NanoString (19). Of the 23 miRNAs comprising our prognostic signature, only 9 are represented in the InterMEL NanoString panel. When our meta-analysis was restricted to the 125 miRNAs included in this panel, these same 9 miRNAs (hsa-miR-451, hsa-miR-21, hsa-miR-200c, hsa-miR-223-3p, hsa-miR-203, hsa-miR-205, hsa-miR-363-3p, hsa-miR-135b-5p, hsa-miR-29a) met the significance thresholds and all showed directionally concordant associations with ulceration in the InterMEL cohort (**Table S4**). The remaining 14 miRNAs from the prognostic signature are not represented in the InterMEL panel and therefore could not be assessed in this cross-validation.

To confirm global transcriptomic reproducibility beyond strict significance cutoffs, Gene Set Enrichment Analysis (GSEA) was performed using InterMEL-defined reference sets. Both the independent upregulated and downregulated miRNA sets were significantly enriched at the exact corresponding extremes of our meta-analysis ranking (adjusted P-value = 0.0001 for upregulated, and 0.028 for downregulated miRNAs), confirming robust cross-platform consensus.

### Integrated functional enrichment identifies OIS escape and stress-adaptive remodeling as the defining programs of melanoma initiation and ulceration

To explore the biological implications of both signatures, we performed a directional Gene Set Analysis propagating miRNA expression signals to their validated target genes, integrating GO-BP and Reactome annotations (see Supplementary Methods). Full enrichment results, including detailed cluster descriptions and sub-pathway annotations, are provided in Supplementary Results S-R1–5 and **Tables S5–S8**.

The diagnostic scenario (15,684 unique target genes) identified three converging biological themes consistent with OIS disruption as a candidate molecular event characterizing early melanomagenesis. This transition is computationally associated with an extensive post-transcriptional reprogramming centered on mRNA metabolism, deadenylation-dependent decay, and 3’UTR-mediated stabilization, compounded by altered SUMOylation processes (**Figure 2A-C**, **Figure 3 [PB2.2-3]**; **Tables S5, S7**). Concurrently, widespread pathway alterations point toward a predicted modulation of the cell cycle G1/S transition and G1/G2 DNA damage checkpoints through the disruption of CDK-dependent signaling cascades (**Figure 2A-C**, **Figure 3 [PB1.1, PB3.3]**), which aligns with the validated targeting of CDK6 within our network (**Figure 4**). Moreover, the enrichment of terms for cellular senescence and its negative regulation provides robust transcriptomic evidence reflecting OIS escape. Crucially, while the signature preserves cellular identity through the exclusive enrichment of melanosome organization and assembly, it highlights an early acquisition of cytoskeletal plasticity and motility potential mediated by altered Rho GTPase signaling and the candidate impairment of the TGF-β/SMAD axis, consistent with the predicted loss of cytostatic tumor-suppressive control in early melanoma development (**Figure 3 [PB3.2-3]**). **Further results are provided as Supplementary Results S-R1-2.**

**Figure 2.**
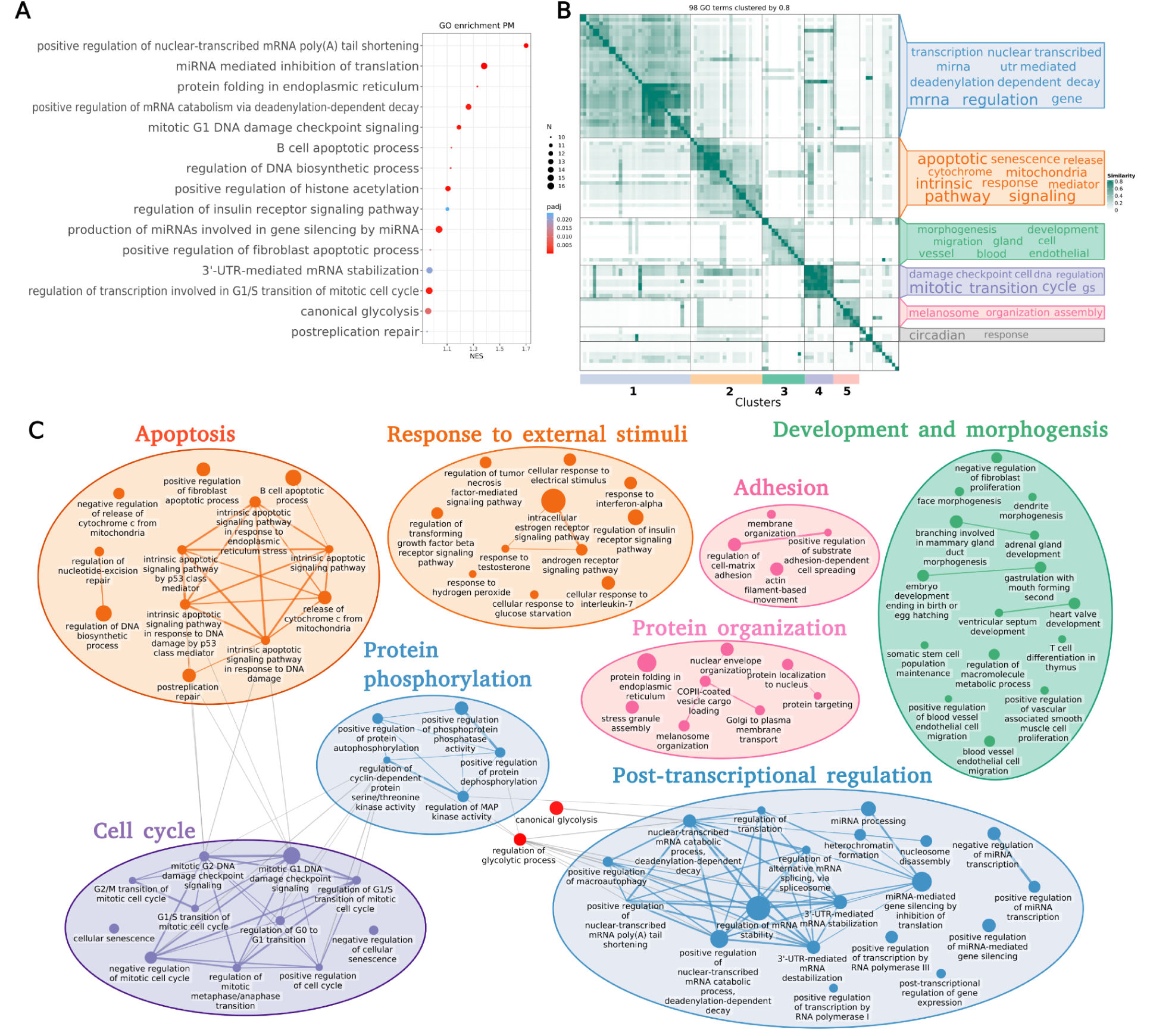
Functional landscape of the diagnostic miRNA signature in primary melanoma. (A) Dot plot illustrating the top fifteen GO-BP processes most significantly enriched in primary melanoma compared to benign nevi. The x-axis represents the Normalized Enrichment Score (NES); dot size and color indicate the gene set size and adjusted P-value, respectively. (B) Semantic similarity matrix of all significant GO-BP terms enriched in primary melanoma. The intensity indicates the degree of semantic similarity. Distinct functional clusters are identified on the axes and summarized by word clouds on the right. (C) Functional network of processes enriched in primary melanoma. Nodes represent GO-BP terms, with size proportional to statistical significance (adjusted P-value). Edges denote gene overlap between terms, with thickness proportional to the degree of similarity. Nodes are colored according to the semantic clusters defined in (B) and grouped into functional modules labeled by their representative biological activity. Only GO-BPs with a LOR cutoff > 0.5 were included.

**Figure 3.**
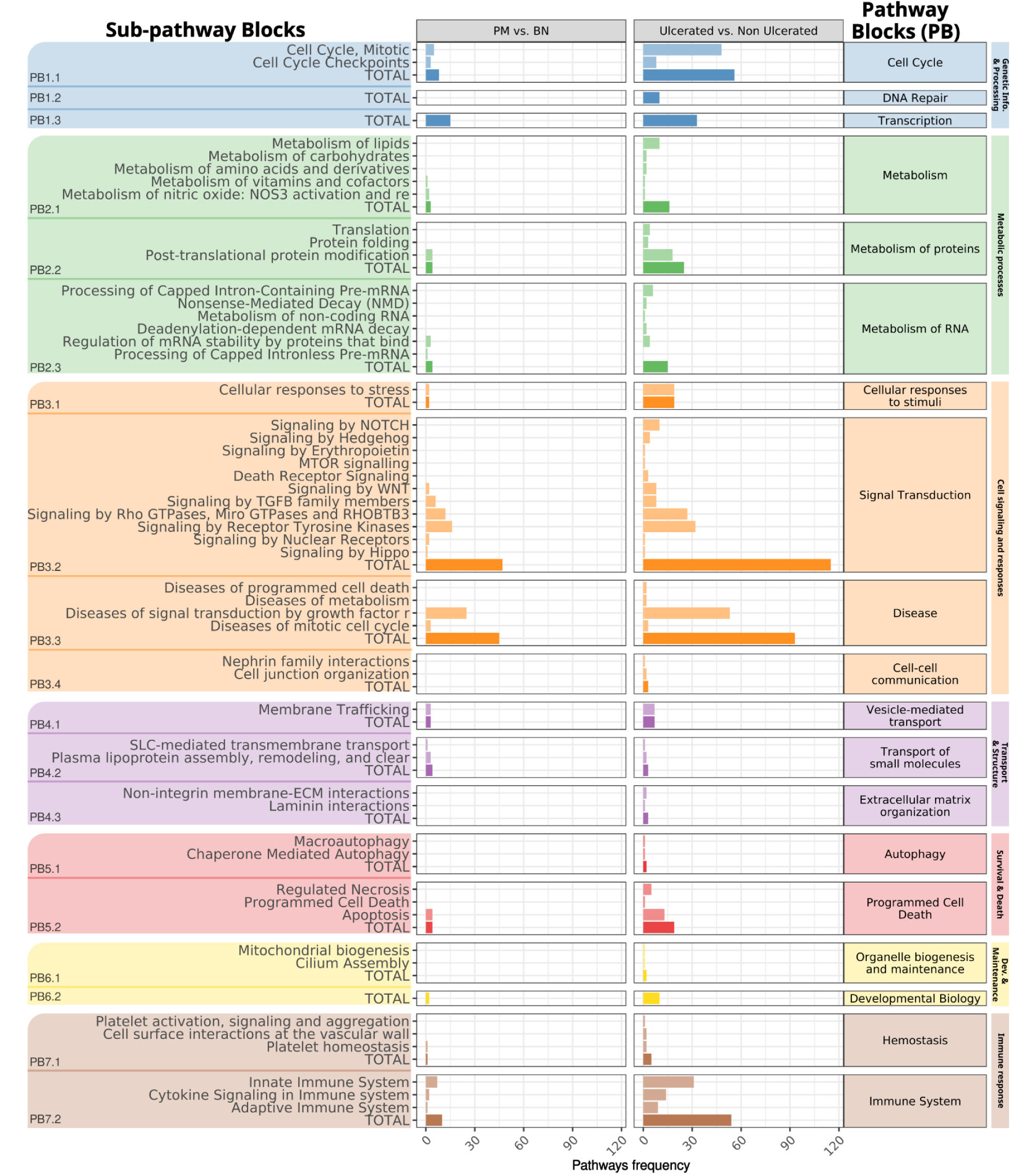
Hierarchical functional pathway signature of the diagnostic and prognostic scenarios. The bar plot displays the frequency distribution of significant Reactome pathways enriched in primary and ulcerated melanoma, categorized into seven distinct **Pathway Blocks (PB)** (right Y-axis, color-coded). Each block groups Top-level pathways, which are further resolved into specific sub-pathways (left Y-axis) detailing precise molecular events. The "TOTAL" bar represents the aggregate count of significant events within each top-level category. The X-axis indicates the number of significant pathways identified in the diagnostic (Primary Melanoma vs. Benign Nevi) and prognostic (Ulcerated vs. Non-Ulcerated) contrasts, allowing for a direct comparison of the functional load associated with tumor initiation versus ulceration-associated risk.

**Figure 4.**
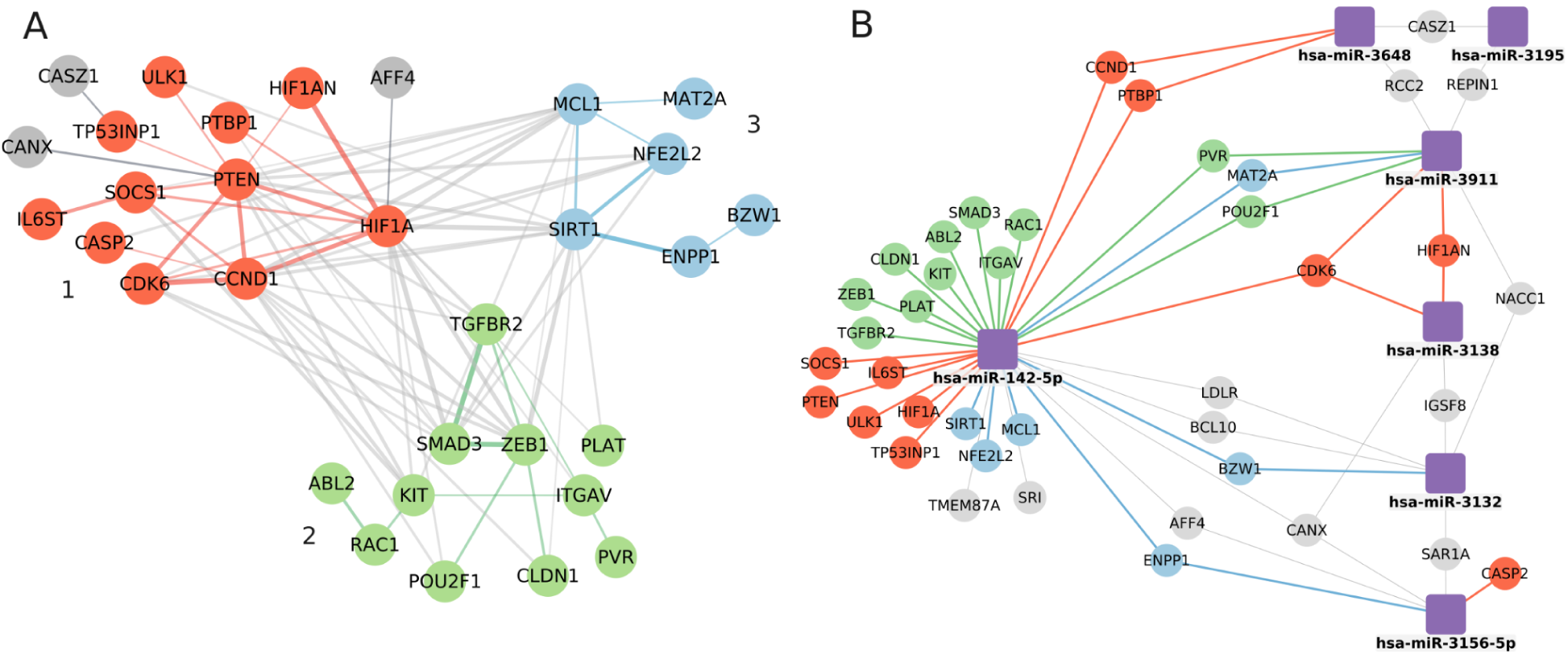
Integrated interaction and regulatory networks linked to early melanomagenesis. (A) Protein-Protein Interaction (PPI) network of validated target genes from miRNAs upregulated in primary melanoma. Nodes represent target genes, while edge thickness indicates the interaction confidence score. The network topology resolves into three distinct dynamic regulatory modules (color-coded): 1) Oncogene-Induced Senescence (OIS) and senescence-associated secretory phenotype (SASP); 2) Proliferation and Epithelial-Mesenchymal Transition (EMT); and 3) Metabolic reprogramming. Unconnected nodes were excluded to highlight the core interactome. (B) miRNA-Target regulatory network. Square nodes represent miRNAs upregulated in primary melanoma, whereas circular nodes correspond to their target genes. Node colors indicate the functional modules defined in (A), while gray nodes refer to targets with lower connectivity or distinct functional roles.

The prognostic scenario (20,150 target genes) identified a broader candidate functional landscape associated with high-risk ulcerated disease. This transition is computationally characterized by extensive structural and pro-invasive remodeling, where alterations in adherens junction assembly and epithelial-mesenchymal transition (EMT) emerged as central, exclusively prognostic processes, alongside dysregulated Rho GTPase and receptor tyrosine kinase (RTK) signaling (**Figure S1A, Figure 3 [PB3.2-4; PB4.3]**; **Tables S6, S8**). Concurrently, widespread pathway alterations and several GO-BP clusters (**Figure S1C**) point toward an active, multidimensional stress-adaptive state. This state is characterized by endoplasmic reticulum (ER) stress, the unfolded protein response, ROS accumulation, and genotoxic stress pathways, including UV radiation, which are compounded by acute proteotoxic stress and translational initiation control (**Figure S1A-C, Figure 3 [PB3.1]**). These adaptive signals are functionally coupled with an expanded programmed cell death module, encompassing autophagy, regulated necrosis, and necroptosis, potentially linking the predicted loss of tissue cohesion with survival cascades such as anoikis resistance (**Figure S1C, Figure 3 [PB5.1-2]**). Finally, the prognostic profile highlights a candidate subversion of the tumor microenvironment, reflected in predicted alterations of modules entirely absent from the diagnostic scenario, such as hemostasis-associated angiogenic signaling, a heavily modulated innate and adaptive immune landscape, along with altered growth factor signaling and developmental processes (**Figure 3 [PB7.1-2; PB3.2-3; PB6.2]**). **Further results are provided as Supplementary Results S-R3-4.**

Comparison of both functional profiles computationally suggests a nested architecture (**Figure 3**). The prognostic signature is predicted to retain core diagnostic alterations while incorporating qualitatively distinct, stress-adaptive modules absent from early transformation. Importantly, shared pathways point toward a candidate functional reorientation: signal transduction is predicted to shift from cytostatic growth factor control (TGF-β, Hippo) toward stress-activated survival cascades (TNF/NF-κB, AKT/PI3K/mTOR), while programmed cell death broadens from intrinsic apoptosis to encompass autophagy and necroptosis. Collectively, these findings are consistent with a progressive model in which early oncogenic checkpoint evasion appears insufficient to account for the high-risk ulcerated phenotype, which instead seems to require an active metabolic and microenvironmental stress adaptation. **Further results are provided as Supplementary Results S-R5.**

### miRNA-driven suppression of senescence and metabolic regulators in primary melanoma

To delineate the candidate regulatory architecture underlying early melanomagenesis, we constructed a skin-tissue-restricted interaction network from validated targets of upregulated diagnostic miRNAs. The resulting Protein-Protein Interaction (PPI) network reveals a highly interconnected core of 31 genes (p = 2.46e-10) topologically segregated into three key predicted modules: OIS/SASP regulation (comprising PTEN, TP53INP1, CDK6, CCND1, CASP2, SOCS1, and IL6ST), cellular proliferation/EMT (integrating TGFBR2, SMAD3, ZEB1, RAC1, ITGAV, KIT, and ABL2), and metabolic reprogramming (centered on SIRT1, HIF1A, MCL1, ENPP1, and NFE2L2) (**Figure 4A**; **Table S9**). This centralized topology suggests that early malignant transformation is computationally associated with coordinated pathway shifts rather than isolated gene events. Crucially, the regulatory network analysis highlights hsa-miR-142-5p as a multifunctional candidate hub (**Figure 4B**), extending its regulation across all three functional modules by simultaneously targeting SIRT1, HIF1A, TGFBR2, RAC1, and CDK6. This convergence positions hsa-miR-142-5p as a critical candidate node integrating metabolic rewiring with senescence evasion and motility. Additionally, hsa-miR-3648, hsa-miR-3138, and hsa-miR-3911 emerge as predicted modulators of the OIS machinery, with hsa-miR-3911 further acting as a potential dual regulator bridging EMT and metabolic control in coordination with hsa-miR-3132 and hsa-miR-3156-5p (**Figure 4B**).

### Upregulated miRNAs in ulcerated melanoma target cell cycle and stress response checkpoints

In the prognostic scenario, the interactome of targets controlled by upregulated miRNAs in ulcerated melanoma unveiled a highly integrated candidate network of 137 genes (p = 1.0e-16), pointing toward a predicted integration of proliferative signaling and surveillance machinery (**Figure 5A**; **Table S10**). The network topology is characterized by prominent connectivity between cell cycle modulators (CDK4, CDK6, E2F1) and sensors of genotoxic or metabolic stress (ATM, HIF1A, PTEN, MDM2), reflecting a coordinated transcriptomic architecture associated with sustained growth under adverse microenvironmental conditions. This network also highlights strong predicted regulation of cell-matrix adhesion, tissue cohesion, and cytoskeletal stability, identifying ICAM1, STMN1, and importantly, RHOB as central candidate nodes, while STAT3 and IL6 function as immune inter-module connectors. The regulatory network (**Figure 5B**) positions hsa-miR-223-3p at the center of this prognostic network due to its high connectivity, directly targeting RHOB, ICAM1, STMN1, and proliferative checkpoints. Additionally, hsa-miR-363-3p is identified as a candidate modulator targeting PTEN and BCL2L11, whereas hsa-miR-196b-5p and hsa-miR-135b-5p are predicted to converge upon a specialized sub-network centering on HIF1A regulation, further reinforcing the stress adaptation model of the high-risk ulcerated phenotype (**Figure 5B**).

**Figure 5.**
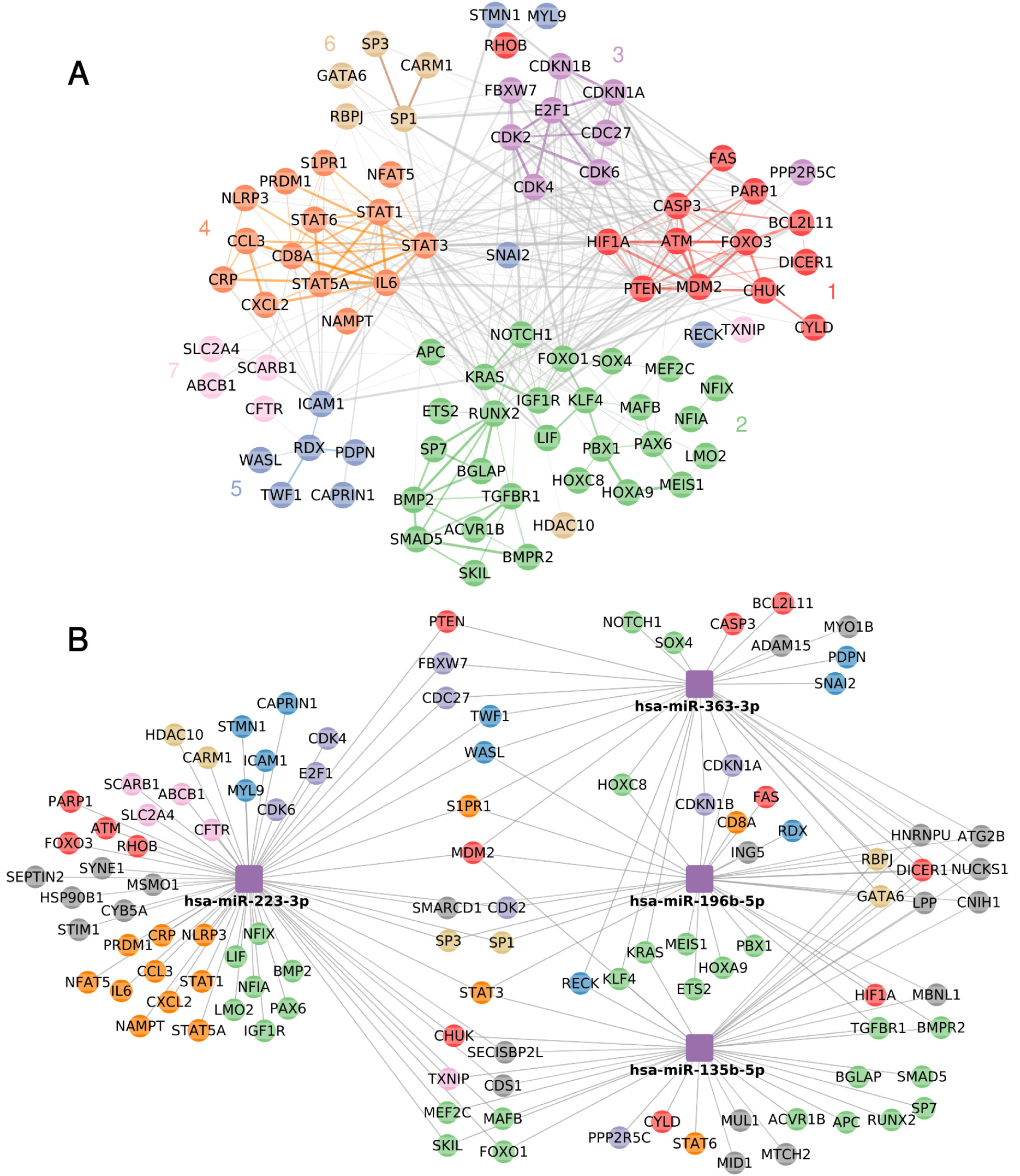
Integrated interaction and regulatory networks characterizing the ulcerated melanoma phenotype. (A) Protein-Protein Interaction (PPI) network of validated target genes of miRNAs upregulated in ulcerated melanoma. Nodes represent target genes, while edge thickness indicates the interaction confidence score. Edges are colored to denote intragroup (matching node color) or intergroup (gray) connections. The network topology resolves into seven distinct dynamic regulatory modules (color-coded): 1) Stress and cell death; 2) Cellular differentiation and proliferation; 3) Cell cycle; 4) Immune system; 5) Mobility and adhesion; 6) Signal transduction and genome maintenance; and 7) Metabolism. Unconnected nodes were excluded. (B) miRNA-Target Regulatory Network. Square nodes represent miRNAs upregulated in ulcerated melanoma, whereas circular nodes correspond to their target genes. Node colors indicate the functional modules defined in (A), while gray nodes refer to targets with lower connectivity or distinct functional roles.

### Ulcerated melanoma is associated with loss of miRNA-mediated repression of growth factor signaling networks

Conversely, targets of downregulated miRNAs in ulcerated melanoma yielded an interconnected network of 50 genes (p = 1.0e-16), highlighting a candidate loss of miRNA-mediated repression over growth factor signaling networks (**Figure S2A**; **Table S11**). The topological organization reveals a dominant predicted core of RTKs and signal transducers, including ERBB2, IGF1R, EGFR, PIK3R2, and SMAD4, interacting with motility and EMT drivers (SNAI2, TWIST1) alongside cell cycle (FOXM1, CDKN1A) and stress response mediators (FOXO4, PDCD4). Regulatory network identifies hsa-miR-489-3p and hsa-miR-499a-5p as highly influential hubs targeting the growth signaling module (**Figure S2B**). Specifically, hsa-miR-489-3p converges with hsa-miR-552-3p, hsa-miR-5010-5p, and hsa-miR-133a-5p to target key growth axes, while hsa-miR-376c-5p is predicted to modulate stress response machinery.

## Discussion

Melanoma progression is conceptually organized around two sequential biological barriers: the escape from Oncogene-Induced Senescence (OIS) during the transition from benign nevi to primary melanoma, and the subsequent adaptation to the hostile, nutrient-deprived ulcerated microenvironment during aggressive tumor progression (20,21). While our meta-analysis cannot establish a direct causal relationship between the diagnostic and prognostic miRNA signatures, which derive from independent patient cohorts, the molecular architecture revealed by each comparison is consistent with each of these two established barriers. Specifically, our data point to hsa-miR-142-5p as a candidate contributor to OIS escape, while the ulceration-associated signature implicates hsa-miR-223-3p in subsequent stress adaptation and invasive remodeling. We interpret these findings within this established two-barrier framework, while acknowledging that longitudinal validation in paired samples across progression stages would be required to confirm a sequential relationship.

The functional profile of the diagnostic signature suggests that dysregulation of G1 checkpoint and DNA repair pathways may compromise the cellular response to DNA damage, potentially through TP53INP1-dependent signaling (22,23). Our functional analysis supports the association of these events with OIS and the senescence-associated secretory phenotype (SASP) (24,25), which may paradoxically shift from tumor-suppressive to tumor-promoting upon chronic activation during melanoma progression (26,27). In this context, network topology positions hsa-miR-142-5p as a candidate central regulator linking OIS escape with malignancy. Although previously described as a prognostic biomarker in several malignancies (28,29), our analysis proposes it may contribute to OIS escape through coordinated modulation of DNA damage response effectors including TP53INP1, alongside metabolic and survival regulators such as SIRT1, HIF1A, and PTEN (30–32).

Mechanistically, OIS depends on stable G1–S arrest mediated by the CDK4/6–RB axis (33,34). Our network predicts that hsa-miR-142-5p and additional OIS-regulatory miRNAs converge on CDK6 and CCND1, potentially weakening G1 checkpoint stability rather than enforcing complete arrest. Notably, hsa-miR-142-5p also links cell-cycle regulation with the SASP by targeting ULK1 and IL6ST. Given that autophagy supports the metabolic demands of SASP (25,35), this miRNA-mediated axis could influence the balance between tumor-suppressive senescence and chronic IL-6-driven inflammation (35,36). Furthermore, the predicted targeting of TGFBR2 and SMAD3 suggests impaired TGF-β-mediated cytostasis (37,38), consistent with the established role of defective TGF-β signaling in melanoma progression and immune evasion (37,39).

Beyond cell cycle modulation, hsa-miR-142-5p may support metabolic plasticity by converging on HIF1A, PTEN, and SIRT1, a node governing glycolipid reprogramming and senescence suppression in melanoma (40–43). Complementing this central axis, miR-205 downregulation, previously characterized as loss of a tumor suppressor constraining proliferation via E2F1 targeting (44), is further linked in our network to altered ZEB1 regulation, consistent with EMT activation (45). Conversely, for the well-established oncogene hsa-miR-21 (46,47), its upregulation within our topology is directly connected to the targeting of PTEN, a key interaction associated with downstream AKT/ERK activation (48). This latter axis connects EMT profiles with metabolic reprogramming, correlating with increased HIF-1α expression and the hypoxic response (49). Collectively, network analysis positions hsa-miR-142-5p as a candidate integrator acting as a "metabolic switch" balancing senescence suppression with survival pathway activation. Notably, the recurrence of hsa-miR-205 and hsa-miR-203 downregulation, alongside hsa-miR-21 upregulation across both comparisons, suggests these markers represent core progression alterations rather than stage-specific events.

Following the initial transformation, tumors with ulceration face severe nutrient deprivation and matrix detachment. While the resulting metabolic crisis and ROS accumulation typically induce anoikis (21,50,51), our functional landscape suggests ulcerated cells could engage HIF1A-associated survival and autophagy pathways to counteract bioenergetic collapse (52–55). Within this context, network topology positions hsa-miR-196b-5p and hsa-miR-135b-5p as candidate modulators of this adaptive reprogramming by targeting HIF1A and the autophagy-related gene ATG2B, highlighting a potential miRNA-driven network linked to anchorage-independent survival.

Reinforcing these adaptations, the predicted loss of repression over EGFR and ERBB2, mediated by downregulation of hsa-miR-133a-5p, hsa-miR-301a-5p, hsa-miR-489-3p, and hsa-miR-552-3p, may sustain MAPK and PI3K/Akt signaling to suppress BCL2L11 and rescue cells from anoikis (50,52,54,56,57). This resistant phenotype is further reinforced by upregulated hsa-miR-363-3p, targeting PTEN and BCL2L11, and hsa-miR-223-3p, which targets TP53INP1, jointly preventing cell death and promoting metastatic potential (30,58,59).

In this adaptive context, the profound downregulation of hsa-miR-200c emerges as a potentially critical event. Its progressive loss across the full spectrum of melanoma progression, along with miR-205, correlates with increased thickness, ulceration, and shorter melanoma-specific survival as an independent prognostic factor (60–62). Mechanistically, while the downregulation of hsa-miR-200c likely reflects the loss of a canonical suppressor constraining ZEB1/2-driven mesenchymal transition and anoikis resistance (63,64),the concurrent predicted loss of hsa-miR-580-3p further derepresses the central EMT inducer TWIST, synergistically reinforcing the pro-invasive remodeling of the ulcerated phenotype (4,21,65). Finally, while general Rho GTPase signaling supports motility, our data highlight a specific predicted mechanism involving RHOB, a motility-limiting tumor suppressor and apoptotic stress sensor, as a key target of hsa-miR-223-3p. The upregulation of hsa-miR-223-3p as an indicator of the ulcerated state is a notable finding, as its regulatory function, documented in myeloid leukemias (66), in melanoma may relieve constraints on invasive behavior, allowing cells to evade stress-induced death (67). Collectively, these network alterations illustrate how miRNA dysregulation coordinates a multi-layered response providing the plasticity required for metastasis.

Several limitations of this study should be considered. First, the mechanistic axes identified, including the miR-142-5p/CDK6/SIRT1 and miR-223-3p/RHOB regulatory modules, represent hypotheses derived from an integrative *in silico* pipeline combining differential expression meta-analysis, database-curated target annotation, and network topology analysis. Each analytical step introduces assumptions that compound across the pipeline. Crucially, while our network construction relies exclusively on previously validated miRNA-target interactions, the biological significance of these candidate axes awaits prospective experimental confirmation. Second, the meta-analysis is based on a limited number of independent datasets per scenario, which inherently introduces platform and processing heterogeneity alongside sample size imbalances. Finally, while we validated the methodological robustness of our prognostic signature against the independent InterMEL dataset (19), differences in sample type, patient populations, and endpoint definitions preclude direct equivalence. Furthermore, no equivalent independent platform exists for the diagnostic comparison, and its robustness therefore rests primarily on the internal consistency of the meta-analytic framework.

In conclusion, our integrative analysis provides a comprehensive map of miRNA alterations associated with melanoma progression, highlighting axes with significant therapeutic potential. Clinically, patients with hsa-miR-142-5p/CDK6 dysregulation might benefit from CDK4/6 inhibitors (e.g., palbociclib), while those exhibiting hsa-miR-223-3p upregulation and EGFR derepression could require combination therapies to overcome BRAF inhibitor resistance (68). To translate these computational predictions into clinically actionable insights, the miR-142-5p/CDK6/SIRT1 axis could be functionally validated in BRAF-mutant melanocyte OIS models measuring senescence markers alongside CDK6 and SIRT1 protein levels. Similarly, the miR-223-3p/RHOB axis could be tested in anchorage-independent growth assays measuring anoikis resistance and RHOB suppression upon miR-223-3p overexpression in non-ulcerated melanoma lines.

## Material and methods

### Dataset selection and search strategy

A systematic review was conducted following Preferred Reporting Items for Systematic Reviews and Meta-Analyses (PRISMA) guidelines (18). We performed a comprehensive search of the Gene Expression Omnibus (GEO) (69) and ArrayExpress (AE) (70) repositories using the keywords: “melanoma”, “microRNAs”, “microRNA”, and “miRNA”. Eligible studies were restricted to publications between 2010-2023, and were filtered based on the following criteria: (i) organism: *Homo sapiens*; (ii) study type: “Non-coding RNA profiling by array" or “expression profiling by array” and (iii) a minimum sample size (n ≥ 12) to ensure adequate statistical power. Conversely, studies were excluded if they met any of the following criteria: (i) lack of focus on melanoma pathology; (ii) absence of miRNA expression data; (iii) use of non-human tissue samples; (iv) experimental designs involving drug treatments or interventions; or (v) use of *in vitro* cell line models, ensuring the analysis focused exclusively on clinical tissue specimens.

### Differential expression and meta-analysis

Raw expression data were independently normalized and quality-controlled using R (71). Intra-study differential expression was modeled using the limma package (72) across two clinical contexts: Primary Melanoma vs. Benign Nevi (diagnostic scenario) and Ulcerated vs. Non-Ulcerated Melanoma (prognostic scenario). The resulting effect sizes (logFC, standard errors, and Benjamini-Hochberg adjusted P-values (BH) (73)) were subsequently combined across studies through a random-effects meta-analysis implemented with the metafor package (74) using the DerSimonian and Laird estimator (75) to weigh inter-study heterogeneity across different platforms. Significance thresholds were set at an adjusted P-value < 0.05 and an absolute logFC > 0.5. **Further methodological details are provided as Supplementary Methods (SM1) along with the integrative bioinformatic workflow in Figure S3.**

### Cross-cohort validation

To evaluate the global transcriptional concordance between our meta-analysis and the InterMEL cohort (19), a Gene Set Enrichment Analysis (GSEA) was performed using the fgsea R package (76). Two independent reference miRNA sets were defined using the InterMEL dataset, comprising miRNAs significantly upregulated and downregulated in association with ulceration. These miRNAs sets were tested against our meta-analytic results ranked by Z-score (logFC/SE). **Further methodological details are provided as Supplementary Methods (SM2).**

### Functional enrichment and Interactome networks

Experimentally validated targets were retrieved from TarBase v8.0 (77) and miRTarBase v9.0 (78). Functional enrichment was performed utilizing a directional logistic regression framework via mdgsa package (79) across GO Biological Process (80) and Reactome annotations (81). A semantic clustering analysis was performed on significant terms (BH < 0.05) using REVIGO (82) and simplifyEnrichment (83) to reduce functional redundancy. The resulting functional networks were visualized in Cytoscape (84).

To construct regulatory interactomes, high-confidence target genes were selected using a strict tripartite filtering strategy in miRTarBase (**Table S12**), prioritizing ’strong evidence’ (luciferase, Western Blot, qPCR) or multi-validated high-throughput targets. Non-physiological interactions were excluded using a tissue-expression filter, retaining only targets with a validated expression score ≥ 2 in skin tissue. Protein-protein interactions and microRNA-target regulatory networks were integrated and visualized using Cytoscape. **Further methodological details are provided as Supplementary Methods (SM3).**

### Data Availability Statement

The data used for the analyses described in this work are publicly available at GEO and AE repositories. The accession numbers of the GEO datasets downloaded are <u>GSE34460</u>, <u>GSE62370</u>, <u>GSE62371</u>, <u>GSE211090</u>, and <u>GSE211092</u>. The accession number of the AE datasets downloaded is <u>E-MTA-B4915</u>. The strategy followed in this systematic review has been described and registered with PROSPERO, and is available via the following link: https://www.crd.york.ac.uk/PROSPERO/view/CRD420261448335.

### Ethics Statement

This research has used anonymised data and metadata from transcriptomic studies available in public repositories. Each of these 6 studies has been conducted in accordance with the ethical guidelines of each institution and country where it was carried out, and is described in detail in each of the reference publications for these studies. *No patients were enrolled during the development of this study*.

## Acknowledgments

The authors thank the Principe Felipe Research Center (CIPF) for providing access to the cluster, co-funded by European Regional Development Funds (FEDER) in the Valencian Community 2014-2020. We would like to thank Dr Patricia P. Centeno of the Cancer Research UK Scotland Institute, for her guidance on this manuscript. The authors also thank Stuart P. Atkinson for reviewing the manuscript. Francisco García García is the guarantor for the work.

This research was supported and partially funded by CIAICO/2023/149 and CIGE/2024/242, from the Consellería de Educación, Cultura y Universidades de la Generalitat Valenciana; by PID2021-124430OA-I00 and PID2023-146836NB-I00 from MICIU/AEI/10.13039/501100011033, and by ERDF/EU. Borja Gómez-Cabañes is supported by a PhD fellowship funded by the Spanish Association Against Cancer in Valencia (PRDVA234163GOME). E.H. is supported by U54CA263001, R01CA274100 and P50CA225450 (National Cancer Institute, NCI/National Institutes of Health, NIH). N.M.E. was supported by F30CA288047 (NCI/NIH).

